# Visual motion computation in recurrent neural networks

**DOI:** 10.1101/099101

**Authors:** Marius Pachitariu, Maneesh Sahani

## Abstract

Populations of neurons in primary visual cortex (V1) transform direct thalamic inputs into a cortical representation which acquires new spatio-temporal properties. One of these properties, motion selectivity, has not been strongly tied to putative neural mechanisms, and its origins remain poorly understood. Here we propose that motion selectivity is acquired through the recurrent mechanisms of a network of strongly connected neurons. We first show that a bank of V1 spatiotemporal receptive fields can be generated accurately by a network which receives only instantaneous inputs from the retina. The temporal structure of the receptive fields is generated by the long timescale dynamics associated with the high magnitude eigenvalues of the recurrent connectivity matrix. When these eigenvalues have complex parts, they generate receptive fields that are inseparable in time and space, such as those tuned to motion direction. We also show that the recurrent connectivity patterns can be learnt directly from the statistics of natural movies using a temporally-asymmetric Hebbian learning rule. Probed with drifting grating stimuli and moving bars, neurons in the model show patterns of responses analogous to those of direction-selective simple cells in primary visual cortex. These computations are enabled by a specific pattern of recurrent connections, that can be tested by combining connectome reconstructions with functional recordings.^*^

**Author summary:** Dynamic visual scenes provide our eyes with enormous quantities of visual information, particularly when the visual scene changes rapidly. Even at modest moving speeds, individual small objects quickly change their location causing single points in the scene to change their luminance equally fast. Furthermore, our own movements through the world add to the velocities of objects relative to our retinas, further increasing the speed at which visual inputs change. How can a biological system process efficiently such vast amounts of information, while keeping track of objects in the scene? Here we formulate and analyze a solution that is enabled by the temporal dynamics of networks of neurons.

## Introduction

One of the most striking properties of biological visual systems is their ability to efficiently cope with the high-bandwidth data streams received from the eyes. To analyze the incoming visual stream, biological systems might employ the efficient coding framework^2–13^. An efficient system would analyze the visual stream and extract its underlying sources, which can often be represented much more concisely. Such sources are, for example, the underlying objects in a visual scene and their instantaneous velocities and directions of motion. We show that such a method can be implemented in a network of interconnected neurons. The model we introduce can qualitatively reproduce known properties of direction-selective simple cells in visual cortex. Furthermore, the required connectivity patterns in the network can be optimized to the statistics of natural movies with a biologically-realistic learning rule. More generally, we propose that a number of time-dependent neural computations could be supported by network dynamics, and might be learned directly from the statistics of the natural world.

### Background

Continuous sequences of images represent complex trajectories through the high-dimensional and nonlinear space of two-dimensional images. Evolutionary pressures on biological visual systems have increased their ability to represent these trajectories efficiently and to distinguish visual motion on a fast time scale. The neural machinery that processes visual motion is therefore well developed^14^, and is assigned in primates to large areas of visual cortex known collectively as the dorsal pathway^15–17^. In this paper we propose a computational model for the very first cortical computation in the dorsal pathway: direction-selectivity as measured in simple cells in primary visual cortex^18^.

The neural mechanisms underlying direction selectivity are still undetermined, but there are experimental indications that it might be accomplished by the temporal dynamics of networks of neurons. For example, recurrent inhibition in the non-preferred direction can account for the direction selectivity of retinal ganglion cells in the retina of rabbits^19^. Unlike orientation and ocular dominance, direction selectivity in cortex requires visual experience to develop^20,21^, perhaps because direction selectivity depends on a specific pattern of lateral connectivity unlike orientation and binocular tuning, which are largely feed-forward and already present at eye-opening. Other experiments have shown that after many exposures to the same moving stimulus, the sequence of spikes triggered in different neurons was also triggered by static stimuli, again suggesting that motion signals in cortex may be generated from lateral connections^22^.

Despite evidence for dynamical processing in cortical networks, the standard model of direction selectivity is phenomenological and describes responses in terms of linear spatiotemporal receptive fields^23–26^. Such models show that the responses of a neuron at time *t* can be predicted by multiplying a spatiotemporal receptive field with the spatiotemporal movie snippet directly preceding time *t*. In some recorded cells, the spatiotemporal receptive fields have a characteristic tilt in the temporal domain, such that they resemble a drifting Gabor stimulus^23,24,26^ (Fig. 1A and supplementary video 1). Out of all stimuli with the same power, such a cell will respond maximally to the drifting Gabor-like stimulus that resembles its receptive field, and respond much less to other directions of movement. Thus that cell is direction selective. Similar models are used in the auditory domain, there called spectrotemporal receptive fields^27^. Although useful for data fitting, receptive field models do not explain the mechanism by which cells compute their responses from the sequences of images presented. Such mechanisms would require knowledge of the stimulus up to a few hundreds milliseconds in the past, yet single neuron timescales are much faster and generally below 10ms^28^. Less well-known models of direction-selectivity incorporate network dynamics as core processes, for example in simulations of cat primary visual cortex^29^. Instead of requiring lagged inputs, such models can maintain information about past stimuli in the internal state of the network, as we show more explicitly below. We expand and generalize this early work to show that a network of neurons with only instantaneous retinal inputs can generate spatiotemporal receptive fields with the required spatiotemporal tilt necessary for selectivity to moving objects. In the rest of the paper we describe learning mechanisms that might lead to such network dynamics in visual cortex. We define mathematically a network which can learn patterns of synaptic connectivity in an unsupervised manner from naturalistic image sequences and show that units in the models have properties similar to those of recorded neurons.

**Figure 1.**
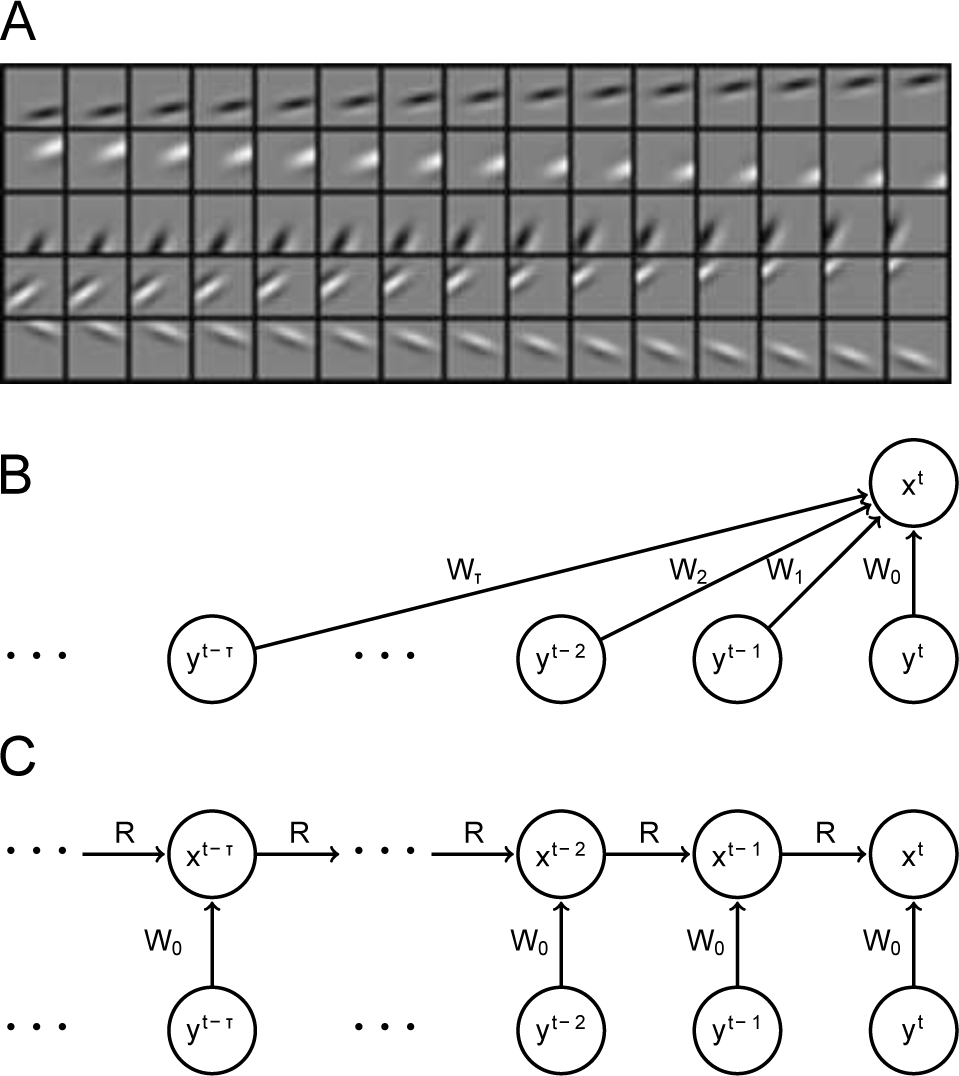
Conversion from a bank of spatio-temporal receptive fields to a recurrent network with instantaneous inputs. (*A*) Five filters out of a bank of 1024 randomly-drawn Gabor spatio-temporal filters. The spatial parameters of the Gabors were chosen randomly from distributions that qualitatively match reported simple cell receptive fields, while the speeds of movements were also drawn independently from a kurtotic distribution with many receptive fields moving relatively slowly, also in agreement with the relative preference of simple cell to slow motions. (*B*) Schematic of the feedforward computation where the variable x_*t*_ represents the output of a simple cell at time t that receives feedforward input from lagged retinal/LGN cells at many previous temporal lags. (*C*) Schematic of a computation performed intrinsically by the network. The activities of a population of neurons x_*t*_ depend only on the instantaneous input, but also on the recent activities of the population x_*t*−1_.

## Results

### Linear neurons with recurrent connectivity enable history-dependent computations

Fig. 1*A*, as well as supplemental video 1, show examples of typical spatiotemporal receptive fields used to model the responses of cortical visual neurons. They consist of an independent filter estimated for each of a number of timelags between a presented stimulus and the neural response. Stimuli presented tens and up to hundreds of milliseconds in the past can affect neural responses. The filters shown in Fig. 1*A* have been obtained from a Gabor function, a well-known parametric model of spatial receptive fields, which has been shifted spatially at constant speeds. They can be used to compute a receptive field's response to an external stimulus in a similar dot-product computation represented graphically in Fig. 1*B.*

Although spatiotemporal filters are useful representations of V1 neuron receptive fields, they have little computational appeal. First, spatiotemporal receptive fields require the specification of a different filter for every timelag in the past, which results in an unnecessarily large number of feedforward connections that have to be learned or specified. Second, the computation requires lagged copies of inputs to be kept available for durations of up to hundreds of milliseconds. Although it has been shown that such lagged inputs are in fact relayed by the LGN^30,31^, lagged LGN neurons still only have a spread of latencies of tens of milliseconds^31^. In addition, storing multiple timelagged retinal images in LGN is greatly hindered by the relatively small number of neurons available there; the LGN is typically thought of as a bottleneck of information transmission^32^.

However, an alternative computation that produces the same output can be devised within biophysically-realistic limits, as represented graphically in Fig. 1*C*. Since neurons are recurrently highly inter-connected in cortex, it is possible to use the dynamics they create to store the relevant information required to estimate the same spatiotemporal patterns. Intuitively, if we discretize time in bins, the activity of neurons at time *t* is available for computations at time *t* + 1. If at time *t* the activity of neurons represents a function of the stimulus many timepoints in the past, by propagation the activity of neurons at time *t* + 1 will also represent the past stimulus. In addition, the instantaneous thalamic input will bring new information about the current spatial stimulus which can be integrated with the old information to produce an estimate of the linear spatiotemporal receptive field. Such a solution to history-dependent computations in visual cortex does not require lagged copies of the input, has many fewer parameters than the purely feedforward filters, can integrate inputs over long periods of time and has reduced computational complexity, in terms of the number of addition/multiplication operations that have to be executed. The appendix gives an approximate calculation of the memory and computation requirements for the feedforward and recurrent filter calculations.

The equations below show a simple derivation that can reparametrize any given set of spatiotemporal basis functions (such as those shown in Fig. 1) to a set of parameters used in a linear recurrent neural network, with instantaneous input filters *W*_0_ and re-current pairwise connections *R*. The reparametrization serves to show a mathematical correspondence between the two solutions to visual history dependence discussed above.

Consider parameterizing a full set of 1024 spatiotemporal receptive fields by their spatial receptive fields (two-dimensional, *n_pix_* by *n_pix_*) at all time lags (a 3rd dimension). We vectorize the two-dimensional filters into a single one-dimensional vector with 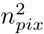 entries and concatenate all filters at time *t* from the entire population of 1024 neurons into a matrix of size 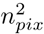 by 1024 and call this matrix *W_t_*. We obtained a sequence of matrices containing the time-lagged filters: *W*_1_, *W*_2_,…*W_t_*…. The linear response of the population of 1024 neurons can be represented by a 1024-dimensional vector *x^t^* that is computed as the following sum

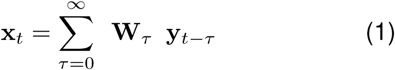

We make the following ansatz, that each of these timelagged matrices *W_t_* can be rewritten in terms of a single recurrent matrix *R* and the initial feedforward matrix at instantaneous timelag *W*_0_:

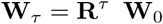

The sum computed in variable *x_t_* in equation 1 can now be rewritten as follows

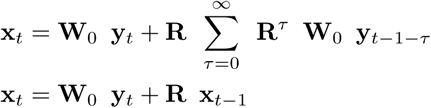

The last equation exactly represents a linear recurrent neural network, with instantaneous feedforward input *W*_0_y_*t*_ and recurrent matrix connectivity **R**. Although we have made the ansatz that each **W**_τ_ can be rewritten as **W**_τ_ = **R**^τ^ **W**_0_, such a matrix **R** may not in general exist. However, under mild assumptions it can be shown that any finite set of sufficiently many spatiotemporal receptive fields can be reparametrized with a matrix **R**. The reparametrization is also easy to obtain under more practical assumptions of a finite and relatively small number of spatiotemporal filters. Under such conditions the reparametrization will not be perfect, but the representation error should be sufficiently small.

We could solve for **R** by linear regression:

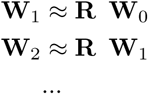

This offers sufficiently precise matrices **R** to reconstruct the entire spatiotemporal profiles of the population at all timelags starting just with the feedforward filter **W**_0_, but it does not maximize the representational capacity of the network, because approximation errors from each independent linear regression add up over each consecutive application of the matrix **R** to the ongoing product **R**^*t*^**W**_*t*−1_. A more exact solution can be obtained by solving the complete linear regression equations

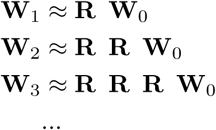

These equations can be (imperfectly) achieved by optimizing the L2-norm of the difference between the left and right hand sides. We performed the optimization using a backpropagation through time fitting procedure^33^.

Solutions **R** to the optimization reconstruct the entire bank of filters very well. Fig. 2*A* and 2*B* show the reconstructed spatial filters for a few selected neurons at timelags of 10 and 20 frames, while the supplemental video 2 shows the full spatiotemporal reconstructions. Note that the reconstruction is perfect at 10 frames in the past, but begins to degrade at 20 frames, with some obvious rippling introduced into the filters. Eventually, the filters at long timelags degrade even more, but they do so in a relatively smooth manner, eventually degrading to 0 because we have penalized the magnitude of the connections *R*. The fits shown in Fig. 2 have been achieved with large *L1* regularization penalties on the parameters, for the purpose of obtaining a sparse matrix *R* with less than 5% overall connectivity, which might more faithfully represent connectivity probabilities for realistic neural networks in cortex.

**Figure 2.**
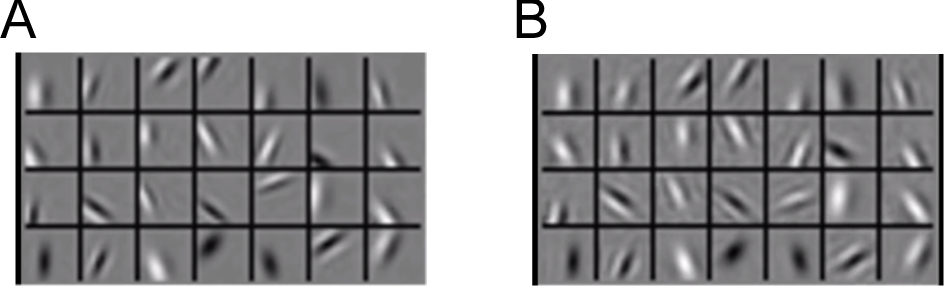
Spatial reconstructions of the bank of filters with intrinsic representations of the recurrent dynamics. (*A*) The reconstructed spatial filters at short timelags (10 frames into the past) are exact. (*B*) The reconstructions at longer timelags (20 frames into the past) are relatively more noisy and begin developing ripples. In general, the quality of the reconstructions is a function of the number of recurrent connections in the network which can store information.

What is the structure of the matrix of pairwise connections *R* determined in this fashion? The eigenvalue spectrum of *R* reveals that most eigenvalues are close to 1, which translate into long timescales of dynamics (Fig. 3*A*). The complex parts of the eigenvalues show that the dynamics are not merely relaxing to 0 over time but in fact gradually shift the stimulus representation from one eigenvector to its pair. Direction-selective simple cells have previously been proposed and implemented with complex-valued basis functions by^11^ specifically to allow responses to rotate across the support vectors provided by the instantaneous receptive field and its π/2 phase-shifted pair. Our implementation provides a representation in terms of real-valued quantities that can be represented by the brain.

**Figure 3.**
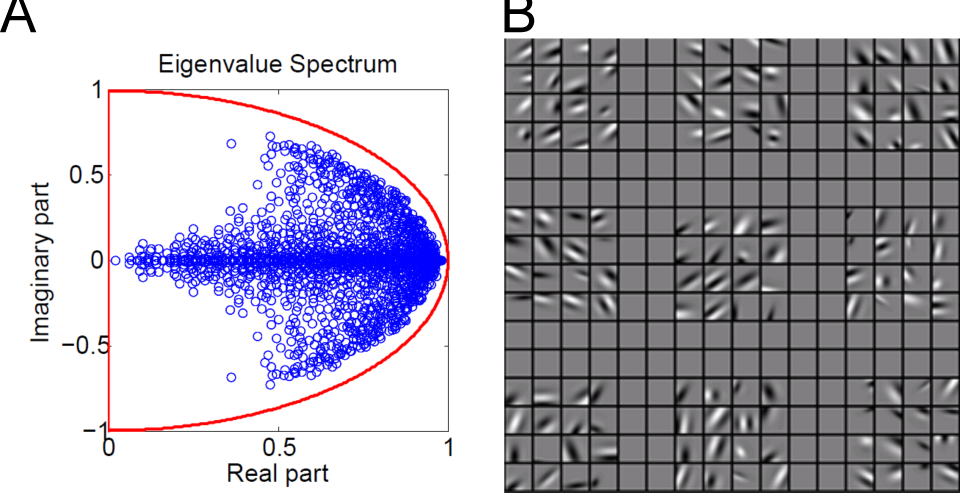
Properties of recurrent connections. (*A*) The eigenvalue decomposition of the recurrent matrix shows a large number of eigenvalues close to 1, which result in long timescales in the spatiotemporal filters. Such long timescales can generate spatial selectivity of a target cell at timelags of 10-20 frames and thus maintain stimulus-information for extended periods of time. (*B*) We can directly inspect what pairs of neurons wire together by looking at their spatiotemporal receptive fields. Neurons with similar prefer directions and orientations wired together preferentially. The figure shows only the instantaneous receptive fields of 9 groups of neurons with strongest connections to 9 target neurons (always shown as the first neuron in the group). All neurons in the same group have similar receptive field properties, and the target receptive field is a linear combination of their receptive fields.

Fig. 3*B* and supplemental video 3 illustrate clusters of neurons with large recurrent connections. Each of the 9 groups shown represents the most strongly-connected neurons to a target randomly chosen neuron (always shown in the top left in the group). Neurons that wire together have similar spatial receptive fields, and they also prefer similar directions of motion (supplemental video 3). Intuitively, a Hebbian learning rule might account for such connectivity patterns in the brain, but it requires modifications. The rest of the paper describes the connectivity patterns that allow visual motion direction estimation. In short, neurons connecting strongly have to prefer similar directions of motion, and their receptive field locations must be spatially aligned to their common preferred direction of motion.

### Selected database of natural videos

We proceed to show that an algorithm based on internal models can adapt a recurrent neural network to the statistics of natural videos via a biologically-plausible learning rule. For naturalistic movies, we selected about 100 short 100 frame long clips from a high resolution BBC wild life documentary (Fig. 4*A*). Clips were chosen only if they seemed on visual inspection to have sufficient motion energy over the 100 frames. The chosen movies were mostly panning shots and close-ups of animals in their natural habitats^†^. Small patches of 16 by 16 pixels were extracted from random positions in the movie clips, along with their temporal sequence in the 100 consecutive frames (Fig. 4*B*).

**Figure 4.**
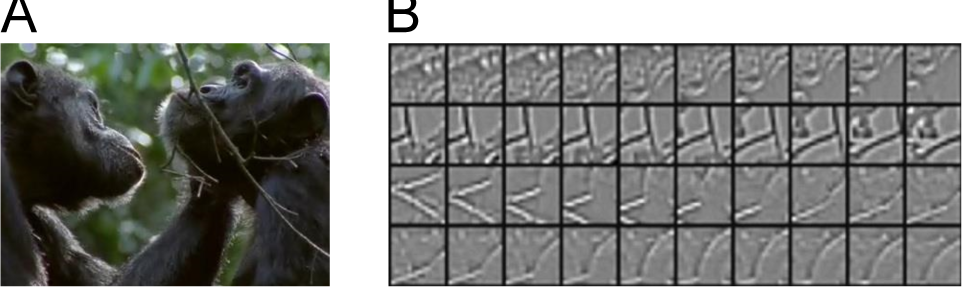
Natural video data used for training the model. (*A*) Snapshot of nature documentary used for learning the statistics of natural movies. (*B*) Four samples of the sequences of patches extracted from the documentary.

The results shown below measure the ability of the model to produce responses similar to those of neurons recorded in primate or cat experiments. The stimuli used in these experiments are typically of two kinds: drifting gratings presented inside circular or square apertures or translating bars of various lengths. These two kinds of stimuli produce very clear motion signals, unlike motion produced by natural movies. In fact, most patches we used in learning contained a wide range of spatial orientations, most of which were not orthogonal to the direction of local translation. We next compare model responses to neural data, and end with an analysis of the connectivity pattern between model neurons that underlies their responses.

### Measuring responses in the model

Unlike neural responses, which are non-negative firing rates, variables in our model could be both positive and negative. We thus separated the positive and negative parts of the Gaussian variables into two distinct sets of responses. This interpretation is relatively common for sparse coding models. Since our inference procedure is deterministic it produces the same response to the same stimulus. We added Gaussian noise to the spatially whitened test image sequences to show robustness of direction selectivity to noise. The amount of noise added was about half the average variance of the stimulus.

### Direction selectivity and speed tuning

Direction selectivity is measured with the following index: DI = 1 - *R*_opp_/*R*_max_. Here *R*_max_ represents the response of a neuron in its preferred direction, while *R*_opp_ is the response in the direction opposite to that preferred. This selectivity index is commonly used to characterize neural data. To define a neuron's preferred direction, we inferred latent coefficients over many repetitions of square gratings drifting in 24 directions, at speeds ranging from 0 to 3 pixels/frame in 0.25 steps. The periodicity of the stimulus was twice the patch size, so that motion locally appeared as an advancing long edge. The neuron’s preferred direction was the direction in which it responded most strongly, averaged over all speeds. Once a preferred direction was established, we defined the neuron’s preferred speed, as the speed at which it responded most strongly in its preferred direction. Finally, at this preferred speed and direction, we calculated the DI of the neuron. Similar results were obtained if we averaged over all speed conditions.

We found that most neurons in the model had good receptive fields and direction-selective responses (Fig. 5*A* and *B*). We cross validated the value of the direction index with a new set of responses to obtain an average DI of 0.65, with many neurons having a DI close to 1 (Fig. 5*C*). 714 of 1024 neurons were classified as direction-selective, on the basis of having DI > 0.5. Distributions of direction indices and optimal speeds show good tuning to moving stimuli (Fig. 5*C*). A neuron’s preferred direction was always close to orthogonal to the axis of its Gabor receptive field, except for a few degenerate cases around the edges of the patch. We defined the population tuning curve as the average of the tuning curves of individual neurons, each aligned by their preferred direction of motion. The DI of the population was 0.66. Neurons were also speed tuned, in that responses could vary greatly and systematically as a function of speed and DI was non-constant as a function of speed (Fig. 5*B*). Usually at low and high speeds the DI was 0, but in between a variety of responses were observed. Speed tuning is also present in recorded V1 neurons^34^, and could form the basis for global motion computation based on the intersection of constraints method^35^.

**Figure 5.**
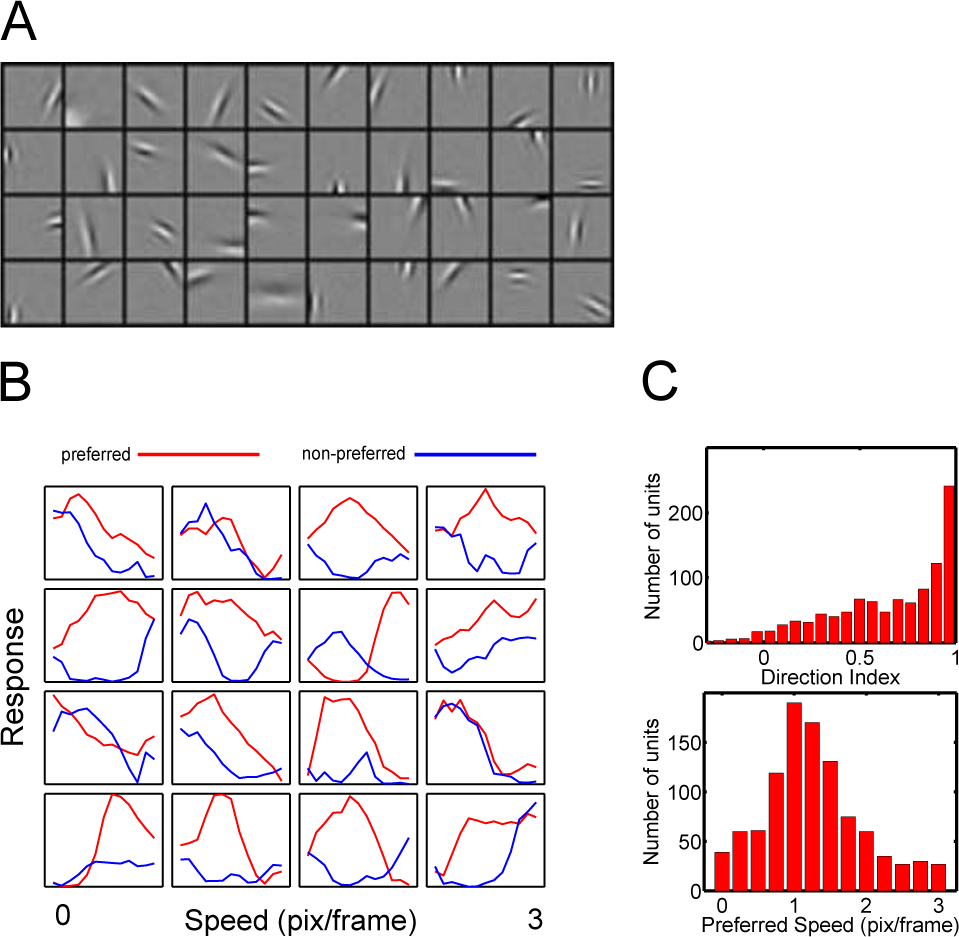
Properties of the learned representations: static receptive fields, speed tuning, direction index and pairwise connectivities. (*A*) Population of static filters derived by the model. (*B*) Speed tuning of 16 randomly chosen neurons. Note that some neurons only respond weakly without motion, some are inhibited in the non-preferred direction compared to static responses and most have a clear peak in the preferred direction at specific speeds. (*C*) top: Histogram of direction selectivity indices. bottom: Histogram of preferred speeds.

### Vector velocity tuning

To get a more detailed description of single-neuron tuning, we investigated responses to different stimulus velocities. Since drifting gratings only contain motion orthogonal to their orientation, we switched to small (1.25pix x 2pix) drifting Gabors for these experiments. We tested the network's behavior with a full set of 24 Gabor orientations, drifting in a full set of 24 directions with speeds ranging from 0.25 pixels/frame to 3 pixels/frame, for a total of 6912 = 24 x 24 x 12 conditions with hundreds of repetitions of each condition. For each neuron we isolated its responses to drifting Gabors of the same orientation travelling at the 12 different speeds in the 24 different directions. We present these for several neurons in polar plots (Fig. 6*A*). Note responses tend to be high to vector velocities lying on a particular line. In the next section we show that these are the so called constraint lines.

**Figure 6.**
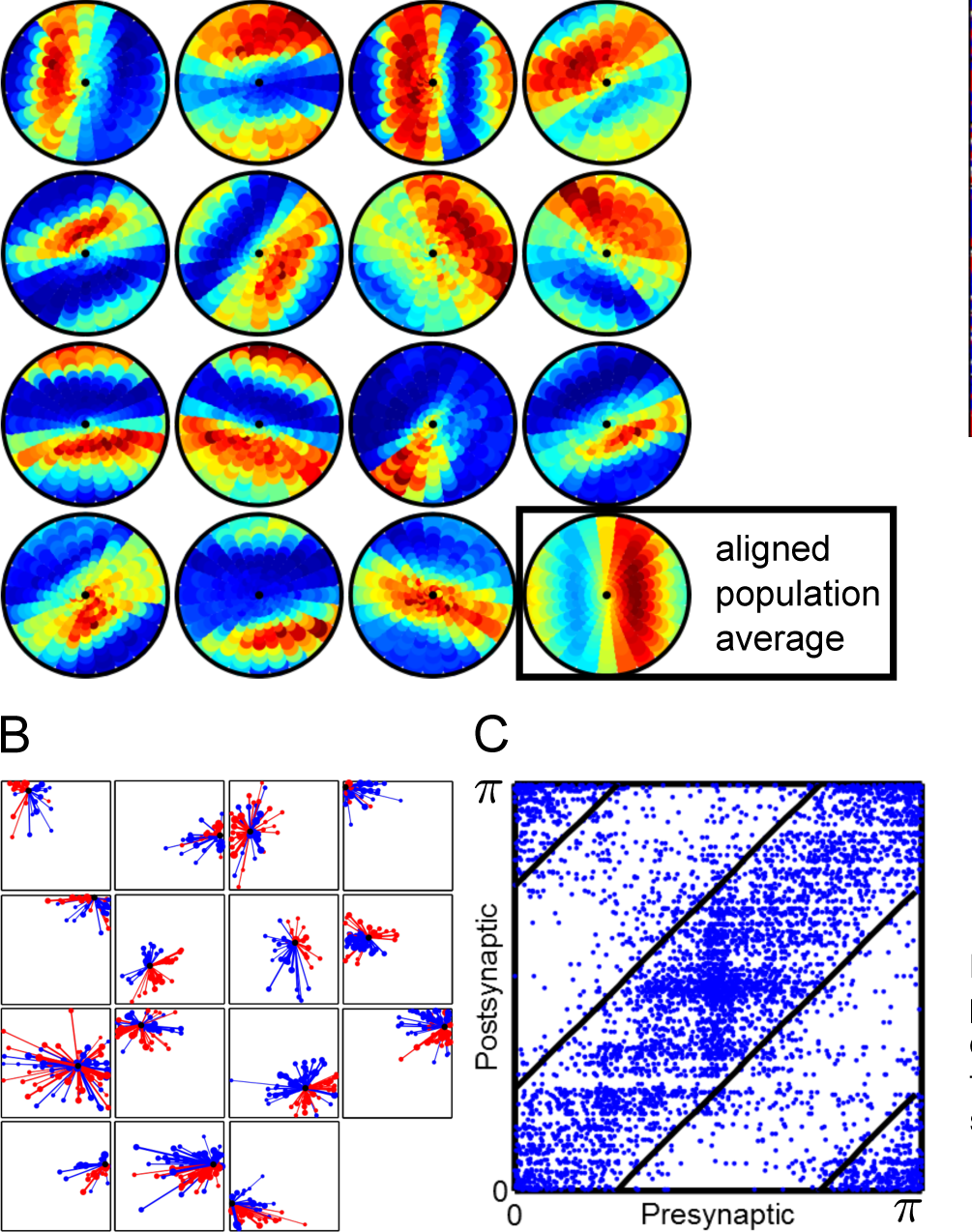
Connectivity patterns between neurons enable selective responses to moving bars. (*A*) The polar plots show the responses of neurons presented to small, drifting Gabors that match their respective orientations. Every disc in every polar plot represents one combination of speed and direction and color represents the magnitude of the response. The vector from the center of the polar plot to the center of each disc is proportional to the vector displacement of each consecutive frame in the stimulus sequence. The last polar plot shows the average of the responses of the entire population, with each neuron aligned by its preferred direction. (*B*) Each plot is based on the outgoing connections of a random subset of direction-selective neurons. The spatial locations of the RFs of root neurons are shown as filled black circles. Filled red/black circles show neurons to which the root neurons have strong positive/negative connections. The width of the connecting lines and the area of the filled circles are proportional to the strength of the connection.(*C*) For each of the 10 strongest excitatory connections per neuron we plot an asterisk indicating the orientation selectivity of pre and post-synaptic units.

### Connectomics in silico

The most obvious connectivity pattern, clearly visible for single neurons (Fig. 6*B*), shows that neurons in the model excite other neurons in their preferred direction and inhibit neurons in the opposite direction. This asymmetric wiring naturally supports direction selectivity in combination with the second pattern described below. The connectivity pattern emerges gradually during learning (Fig. 7*A*, *B* and *C* and supplemental video 4).

**Figure 7.**
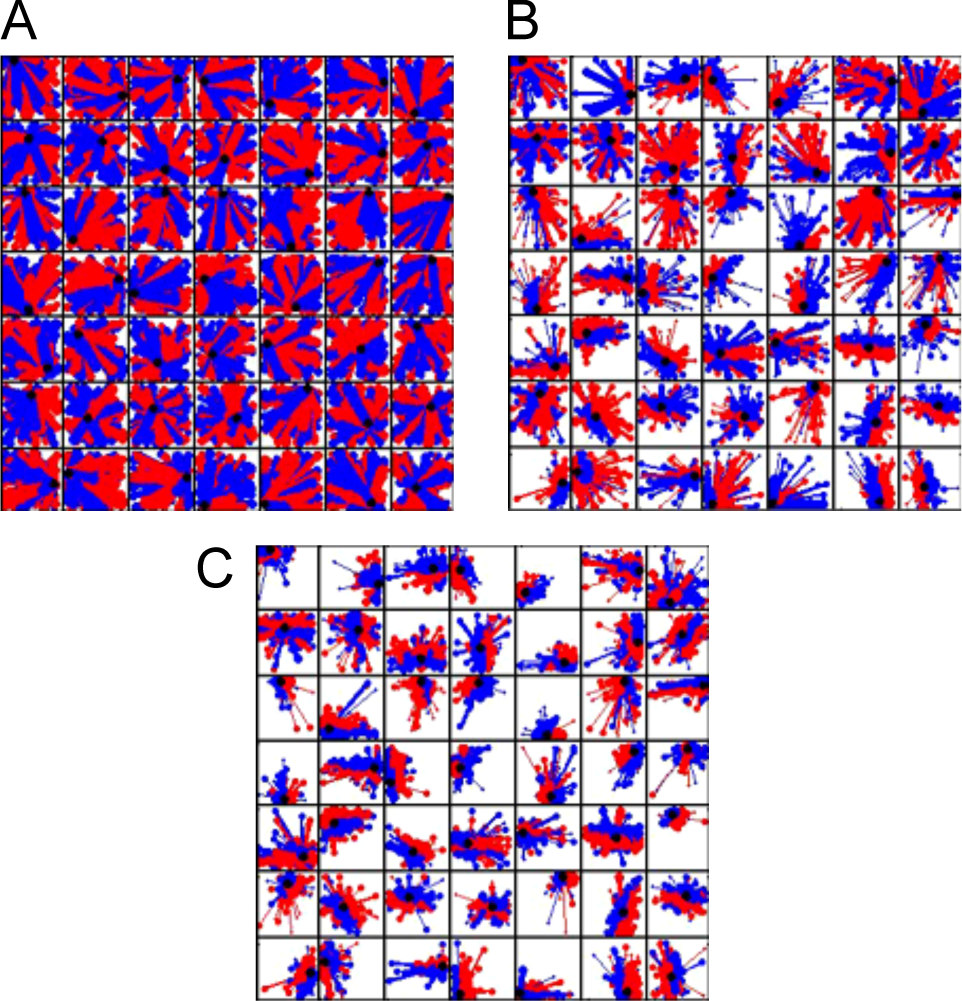
Connectivity changes during learning (see also supplemental video 4). (*A*) We initialized the network with random connections between neurons. (*B*, *C*) During learning, connections gradually sparsify and become asymmetric between the two sides of the receptive fields of each cell.

Asymmetry is not sufficient for direction selectivity to emerge. In addition, strong excitatory projections have to connect together neurons with similar preferred orientations and similar preferred directions. Only then will direction information propagate in the network in the identities of the active neurons. We considered for each neuron its 10 strongest excitatory outputs and calculated the expected deviation between the orientation of these outputs and the orientation of the root neuron. The average deviation was 23°, half the expected deviation if connections were random (Fig. 6*C*). The same pattern held when we considered the strongest excitatory inputs to a given neuron with an expected deviation of orientations of 24°. We could not directly measure if neurons connected together according to direction selectivity because of the sign ambiguity of **x**^*t*^ variables. Neurons connected asymmetrically with respect to their RF axis (Fig. 6*B*), and they also responded to motion primarily in the direction orthogonal to their connectivity axis (Fig. 6*A*). Since direction tuning is a measure of the *incoming* connections to the neuron, we can qualitatively assess that recurrence primarily connected together neurons with similar direction preferences.

We also observed that neurons mostly projected strong excitatory outputs to other neurons that were aligned parallel to the root neuron’s main axis (Fig. 6*B*)). This reminds of the aperture problem: locally all edges appear to translate parallel to themselves. A neuron *X* with a preferred direction *v* and preferred speed *s* has a so-called constraint line (CL), parallel to the Gabor’s axis. When the neuron is activated by an edge *E*, the constraint line is formed by all possible future locations of edge *E* that are consistent with global motion in the direction *v* with speed *s*. Due to the presence of long contours in natural scenes, the activation of *X* can predict at the next time step the activations of other neurons with RFs aligned on the CL. Our likelihood function encourages the model to learn to make such predictions as well as it can, and it discovers the constraint line solution. To quantify the degree to which connections were made along a CL, we fit for each neuron a 2D Gaussian to the distribution of RF positions of its 20 strongest-connected neurons, each further weighted by its strength (the filled red circles Fig. 6*B*). The major axes of the fitted Gaussians were aligned to the root neuron axis by less than 15° in 862 out of 1024 neurons.

Perhaps the strongest manifestation of the CL tuning property of neurons in the model can be seen in their responses to small stimuli drifting with different vector velocities. Many of the neurons in the model responded best when the velocity vector ends on the constraint line (Fig. 6*A*) and a similar pattern held for the aligned population average.

It is known that neurons in V1 are more likely to connect with neurons of similar preferred orientations situated as far as 4 mm / 4-8 minicolumns away^36^. The model presented here makes further predictions: that outgoing connections are made to neurons with similar preferred directions and with receptive field centers displaced along the preferred direction, and located along a constraint line parallel to the receptive field axis.

## Discussion

### Sequence learning

Mechanisms for learning connectivity patterns between neurons have been proposed and found in the form of spike timing-dependent plasticity (STDP). Specific models incorporating STDP treat visual motion as a general sequence learning problem^37,38^. Visual motion processing can thus be seen as a special case of the general problem of sequence learning^39^. Many structures in the brain show various forms of sequence learning, and recurrent networks of neurons can naturally produce learned sequences through their dynamics^40,41^. It has been suggested that the reproduction of remembered sequences in the hippocampus has an important navigational role. Similarly, motor systems are able to generate the sequences of control signals that drive appropriate muscle activity. Thus many neural sequence models are fundamentally generative. In contrast, it is not evident that V1 should need to reproduce the learnt sequences of retinal input that represent visual motion, although it has been recently shown that they do^22^. While generative modelling provides a powerful mathematical device for the construction of inferential sensory representations, the role of actual generation has been debated. Is there really a potential connection then, to the generative sequence reproduction models developed for other areas?

One possible role for explicit sequence generation in a sensory system is for prediction. Predictive coding has been proposed as a central mechanism to visual processing^42^ and even as a more general theory of cortical responses^43^. More specifically as a visual motion learning mechanism, sequence learning forms the basis of an earlier simple toy but biophysically realistic model based on spike-time dependent plasticity (STDP) at the lateral synapses of a recurrently connected network^38^. Fig. 8 shows a schematic of this earlier model which describes one of the basic mechanisms we incorporate as well. Despite its intuitive properties, this early STDP-based model lacks a functional interpretation or a computational objective, that might allow it to scale up in size and be able to compute in natural environments. Predictive properties emerge in networks with the relevant STDP learning rule, but they do not directly optimize a well-defined functional objective. We were able to generalize the neural network predictive model to natural video by reformulating the early models of sequence learning within a generative framework. The framework specifies a well-defined functional objective and allows us to derive learning rules at the lateral connections of the network that directly optimize this objective. The functional objective we derive directly specifies that the model should be adapted to the spatiotemporal statistics of natural videos. Unlike the direct predictive-coding approach, a generative model requires an internal model of the visual world that can be used to make inferences about the current state of the world, and predictions about future states. Thus, such a system can function as a predictor, but does so within a structured model that needs to be adapted to the statistics of the environment.

**Figure 8.**
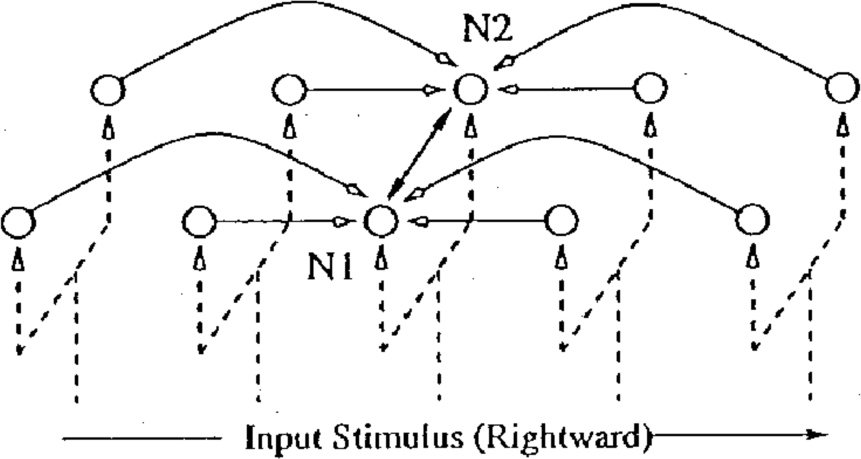
Toy sequence learning model with biophysically realistic neurons from^38^. Neurons N1 and N2 have the same RF as indicated by the dotted line, but after STDP learning of the recurrent connections with other neurons in the chain, N1 and N2 learn to fire only for rightward and respectively leftward motion.

**Figure 9.**
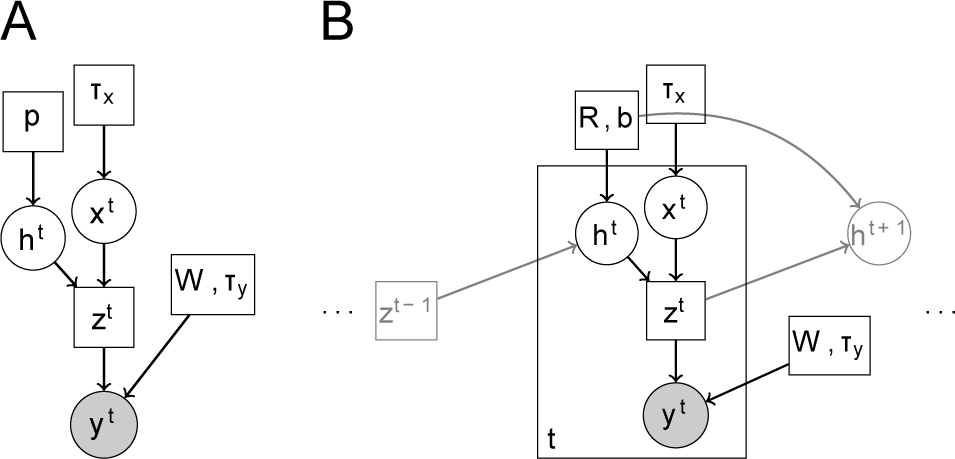
Graphical models used — the relationships between variables (neural activities, images/retinal activities) are shown and the parameters describe mean activities and connectivity parameters. (*A*) Graphical model representation of the generative model of still images. (*B*) Graphical model representation of the bgG-RNN with two consecutive time slices. The square box represents that the variable **z**_*t*_ is not random, but is given by **z**_*t*_ = **x**_*t*_ ○ **h**_*t*_.

Generative models of static natural visual scenes have a long history in computational neuroscience^8,9,44–46^. They have been used to argue that learning and inference in the brain are adapted to the natural environments an animal inhabits. Organisms use an internal model of the external world to decode information from the senses and translate that information into an abstract format more suitable to be used in behavioral programs. An adapted brain would detect the underlying causes of pixel movements on the retina: specific objects moving in the world and/or self-induced optic flow. Such decoding of retinal information requires an accurate and well-adapted internal model. Do brains adapt to the natural world they are embedded in? Biologically, it is well-established that properties of cortical neuron responses change depending on the statistics of experienced visual sensations^47–52^, so it does seem like the internal model is continually-changed based on the statistics of the world.

### Previous models

Many previous low-level generative models of image sequences have treated time as a third dimension in a sparse coding problem^12^. These approaches have thus far failed to map to neural architecture as they have been implemented with noncausal computations. Furthermore, the spatiotemporal sensitivity of each learned "neuron" is determined by a separate three-dimensional basis function, requiring very many variables to encode all possible orientations, directions of motion and speeds. Cortical architecture points to a more distributed formation of motion representation, with temporal sensitivity determined by the interaction of neurons with different spatial receptive fields.

Other models of visual motion analyze the slowly-changing features of the visual input and propose that complex cells are slow feature learners^46,53^. However, these models are not expressive enough to encode visual motion and are more specifically designed to encode image dimensions invariant in time. As such they in fact discard aspects of video that change in time, such as motion.

### History-dependent neural computations through learnt recurrent dynamics

We have shown that a network of recurrently-connected neurons can learn to discriminate motion direction at the level of individual neurons. The model neurons can be interpreted as predicting the motion of the stimulus from the lateral inputs they receive. These inputs are however not sufficient in themselves to produce a response; the prediction also has to be consistent with the bottom-up input. Thus, we see a number of reasons to propose that direction selectivity in cortex may develop and be computed through a mechanism analogous to the one we have developed here. Recent experiments^54^ have shown that neurons with similar functional responses wire together preferentially, and that this pattern of connectivity is not present at eye opening^55^, but emerges afterwards^20,21^. However, it remains to be shown whether neurons tuned to similar directions of motion wire together preferentially and with the spatial relationships we have described.

## Methods

### Probabilistic Recurrent Neural Networks

In this section we introduce the binary-gated Gaussian recurrent neural network as a generative model of sequences of images. The model belongs to the class of nonlinear dynamical systems. Inference methods in such models typically require expensive variational^56^ or sampling based approximations^57^, but we found that a low cost online filtering method works sufficiently well to learn an interesting model. We begin with a description of binary-gated Gaussian sparse coding for still images and then describe how to define the dependencies in time between variables.

### Binary Gated Gaussian Sparse Coding (bgG-SC)

Binary-gated Gaussian sparse coding (also called spike-and-slab sparse coding^58‡^) may be seen as a limit of sparse coding with a mixture of Gaussians priors^13^ where one mixture component has zero variance. Mathematically, the data **y**^*t*^ is obtained by multiplying together a matrix **W** of basis filters with a vector **h**^*t*^ ○ **x**^*t*^, where ○ denotes the operation of Hadamard or element-wise product, **x**^*t*^ ∈ ℝ^*N*^ is Gaussian and spherically distributed with standard deviation *τ_x_* and **h**^*t*^ ∈ {0,1}^*N*^ is a vector of independent Bernoulli-distributed elements with success probabilities **p**. Finally, small amounts of isotropic Gaussian noise with standard deviation *τ_y_* are added to produce **y**^*t*^.

For notational consistency with the dynamic version of this model, the *t* superscript indexes time. The joint log-likelihood is

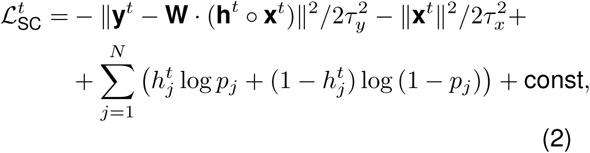

where *N* is the number of basis filters in the model. By using appropriately small activation probabilities **p**, the effective prior on **h**^*t*^ ○ **x**^*t*^ can be made arbitrarily sparse. Probabilistic inference in sparse coding is intractable but efficient variational approximation methods exist. We use a very fast approximation to MAP inference, the matching pursuit algorithm (MP)^59^. Instead of using MP to extract a fixed number of coefficients per patch as usual, we extract coefficients for as long as the joint log-likelihood increases. Patches with more complicated structure will naturally require more coefficients to code. Once values for **x**^*t*^ and **h**^*t*^ are filled in, the gradient of the joint log likelihood with respect to the parameters is easy to derive. Note that 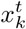 for which 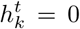 can be integrated out in the likelihood, as they receive no contribution from the data term in (2). Due to the MAP approximation, only the **W** can be learned. Therefore, we set 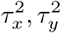 to reasonable values, both on the order of the data variance. We also adapted *p_k_* during learning so that each filter was selected by the MP process a roughly equal number of times. This helped to stabilise learning, avoiding a tendency to very unequal convergence rates otherwise encountered.

When applied to whitened small patches from images, the algorithm produced localized Gabor-like receptive fields as usual for sparse coding, with a range of frequencies, phases, widths and aspect ratios. We found that when we varied the average number of coefficients recruited per image, the receptive fields of the learned filters varied in size. For example with only one coefficient per image, a large number of filters represented edges extending from one end of the patch to the other. With a large number of coefficients, the filters concentrated their mass around just a few pixels. With even more coefficients, the learned filters gradually became Fourier-like.

During learning, we gradually adapted the average activation of each variable 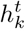 by changing the prior activation probabilities *p_k_*. For 16x16 patches in a twice overcomplete SC model (number of filters = twice the number of pixels), we found that learning with 10-50 coefficients on average prevented the filters from becoming too much or too little localized in space.

## Supporting Information

**S1 Video. Supplemental movie 1.**

**S2 Video. Supplemental movie 2.**

**S3 Video. Supplemental movie 3.**

**S4 Video. Supplemental movie 4.**

## S1 Appendix. Complexity order for recurrent networks versus feedforward filters.

Floating point operations per seconds (FLOPS) are an objective measure of the efficiency of an algorithm. We use a similar measure to estimate the efficiency of computing spatiotemporal neural responses from spatiotemporal visual stimuli. Matters of efficiency are extremely important in vision. The raw amount of visual information available to the senses far exceeds even the computational capacity of the brain and only by selective attention (not discussed in this paper) can it be well encoded by the brain. Neurons in the fovea and up to 5 degrees of visual field represent less than one degree portions of visual space and many columns of neurons with all the required computational apparatus need be replicated throughout striate visual cortex to cover up the entire visual space. Direction-selectivity is just one of the many functions of V1 neurons, with orientation selectivity, color processing and space invariance (complex cells) being equally important qualities of images.

Suppose the spatiotemporal filters are *l_x_* by *l_y_* by *n_t_* in size (space by space by time, a typical example would be 12 by 12 by 30). A full V1 column may employ a number on the order of *N* = 1024 spatiotemporal filters. The number of feed-forward operations necessary to compute the dot-product between the filters and the feedfoward input is thus 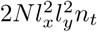. In contrast, the number of recurrent operations necessary for the same computation is 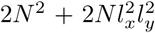. The second term in this sum is smaller than the number of feedforward operations by a factor of *n_t_*, which could be up to 30 (assuming 5ms bins and history-dependence for up to 150ms in the past). The first term may in principle be large, but as discussed above recurrent connectivity in cortex is sparse, hence the total number of non-zero connections in the matrix **R** may be a small fraction of all possible connections, reducing 2*N*^2^ to 2*pN*^2^ with *p* the probability of connection between all pairs of neurons. Sparse connectivity does not impair the quality of the representation, because only neurons with similar spatio-temporal receptive fields need to be wired together.

We see thus that the computational complexity of recurrent neural networks is much reduced compared to simple linear spatiotemporal filters. This advantage mimics the advantage offered by so-called IIR (infinite impulse response) filters over FIR (finite impulse response) filters well-known in the signal processing literature.

The second efficiency concern to any algorithm is its memory requirement. Modern computational hardware indeed improves primarily by making memory access more efficient, and the brain's parallel computational hardware has often been thought to be efficient mostly through its distributed memory systems and the bandwidth of information that travels within and across brain areas continuously. A quick calculation shows that that the memory requirements of recurrent networks are straightforwardly *n_t_* times less than feedforward algorithms, simply because lagged copies of stimulus inputs do not need to be stored at all.

Another algorithmic advantage to the recurrent network is its relatively small number of parameters. If these parameters are indeed learned or optimized to the statistics of the natural world, then an algorithm with fewest free parameters would be optimized the quickest and most efficiently, and would be able to generalize better to new never-before-seen images and sequences-of-images.

Given the significant computational advantages of using a recurrent network to compute visual motion properties, it would be surprising if such a solution was not implemented in neural circuitry. After all, the brain already has the necessary hardware to implement it (the recurrent connections).

## S2 Appendix. Binary-Gated Gaussian Recurrent Neural Network (bgG-RNN).

To obtain a dynamic hidden model for sequences of images {**y**^*t*^} we specify the following conditional probabilities between hidden chains of variables **h**^*t*^, **x**^*t*^:

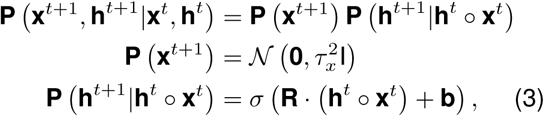

where **R** is a matrix of recurrent connections, **b** is a vector of biases and *σ* is the standard sigmoid function *σ*(*a*) = 1/(1 + exp(−*a*)). Note how the **x**^*t*^ are always drawn independently while the conditional probability for **h**^*t*+1^ depends only on **h**^*t*^ ○ **x**^*t*^. We arrived at these designs based on a few observations. First, similar to inference in SC, the conditional dependence on **h**^*t*^ ○ **x**^*t*^, allows us to integrate out variables **x**^*t*^, **x**^*t*+1^ for which the respective gates in **h**^*t*^, **h**^*t*+1^ are 0. Second, we observed that adding Gaussian linear dependencies between **x**^*t*+1^ and **x**^*t*^ ○ **h**^*t*^ did not modify qualitatively the results reported here. However, dropping **P**(**h**^*t*+1^|**h**^*t*^ ○ **x**^*t*^ in favor of **P**(**x**^*t*+1^|**h**^*t*^ ○ **x**^*t*^ resulted in a model which could no longer learn a direction-selective representation. For simplicity we chose the minimal model specified by (3). The full log likelihood for the bgG-RNN is 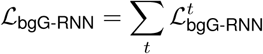 where

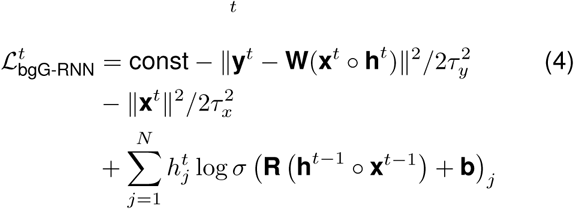

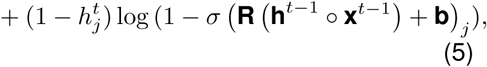

where **x**^0^ = **0** and **h**^0^ = **0** are both defined to be vectors of zeros.

## S3 Appendix. Inference and learning of bgG-RNNs

The goal of inference is to set the values of 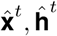 for all *t* in such a way as to minimize the objective set by (5). Assuming we have already set 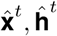 for *t* = 1 to *T*, we propose to obtain 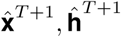 exclusively from 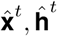. This scheme might be called greedy filtering. In greedy filtering, inference is causal and Markov with respect to time. At step *T* + 1 we only need to solve a simple SC problem given by the slice 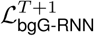 of the likelihood (5), where **x**^*T*^, **h**^*T*^ have been replaced with the estimates 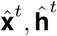. The greedy filtering algorithm proposed here scales linearly with the number of time steps considered and is well suited for online inference. The algorithm might not produce very accurate estimates of the global MAP settings of the hidden variables, but we found it was sufficient for learning a complex bgG-RNN model. In addition, its simplicity coupled with the fast MP algorithm in each 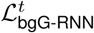 slice, resulted in very fast inference and consequently fast learning. In most scenarios we learned models in under 30 minutes on a quad core workstation.

Due to our approximate inference scheme, some parameters in the model had to be set manually. These are 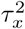 and 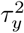 which control the relative strengths in the likelihood of three terms: the data likelihood, the smallness prior on the Gaussian variables and the interaction between sets of **x**^*t*^, **h**^*t*^ consecutive in time. In our experiments we set 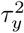 equal to the data variance and 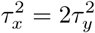. We found that such large levels of expected observation noise were necessary to drive robust learning in **R**.

For learning we initialized parameters randomly to small values and first learned **W** exclusively. Once the filters converge, we turn on learning for **R**. **W** does not change very much beyond this point. We found learning of **R** was sensitive to learning rate. We set the learning rate to 0.05 per batch, used a momentum term of 0.75 and batches of 30 sets of 100 frame sequences. We whitened images with a center-surround filter and normalized the mean and variance of whitened pixel values in the training images to 0 and 1 respectively.

Gradients required for learning *R* show similarities to the STDP learning rule used in^37^ and^38^.

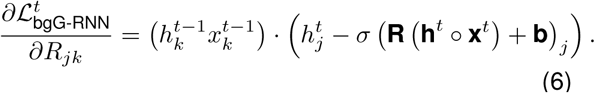

We will assume for neural interpretation that the positive and negative values of **x**^*t*^ ○ **h**^*t*^ are encoded by different neurons. If for a given neuron 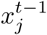 is always positive, then the gradient (6) is only positive when 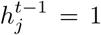 and 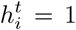 and negative when 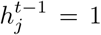 and 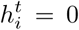. In other words, the connection *R_ij_* is strengthened when neuron *j* appears to cause neuron *k* to activate and inhibited if neuron *j* fails to activate neuron *k*. A similar effect can be observed for the negative part of 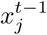. This kind of Hebbian rule is widespread in cortex for long term learning and is used in previous computational models of neural sequence learning that partly motivated our work^38^.

## Acknowledgments

This work was supported by the Gatsby Charitable Foundation. We thank Carsen Stringer for providing comments on the manuscript.

## Footnotes

* Portions of this work have been previously presented^1^

† The closer the camera is to a moving object, the faster it appears to move

‡ Although less evocative, we prefer the term ‘binary-gated Gaussian’ to ‘spike-and-slab’ partly because our slab is really more of a hump, and partly because the spike refers to a feature seen only in the density of the product *h_i_x_i_*, rather than in either the values or distributions of the component variables.

## References

1. Pachitariu, M. & Sahani, M. Learning visual motion in recurrent neural networks. In Advances in Neural Information Processing Systems, 1322–1330 (2012).

2. Doya, K. Bayesian brain: Probabilistic approaches to neural coding (MIT press, 2007).

3. Olshausen, B. A. et al. Emergence of simple-cell receptive field properties by learning a sparse code for natural images. Nature 381, 607–609 (1996).

4. Simoncelli, E. P. & Olshausen, B. A. Natural image statistics and neural representation. Annual review of neuroscience 24, 1193–1216 (2001).

5. Schwartz, O. & Simoncelli, E. P. Natural signal statistics and sensory gain control. Nature neuroscience 4, 819–825 (2001).

6. Karklin, Y. & Lewicki, M. S. Emergence of complex cell properties by learning to generalize in natural scenes. Nature 457, 83–86 (2009).

7. Lewicki, M. S. Efficient coding of natural sounds. Nature neuroscience 5, 356–363 (2002).

8. Karklin, Y. & Lewicki, M. S. A hierarchical bayesian model for learning nonlinear statistical regularities in nonstationary natural signals. Neural Computation 17, 397–423 (2005).

9. Körding, K. P., Kayser, C., Einhäuser, W. & König, P. How are complex cell properties adapted to the statistics of natural stimuli? Journal of neurophysiology 91, 206–212 (2004).

10. Berkes, P., Turner, R. E. & Sahani, M. A structured model of video reproduces primary visual cortical organisation. PLoS computational biology 5, e1000495 (2009).

11. Cadieu, C. & Olshausen, B. Learning transformational invariants from natural movies. Advances in Neural Information Processing 21, 209–216 (2009).

12. Olshausen, B. Learning sparse, overcomplete representations of time-varying natural images. IEEE International Conference on Image Processing (2003).

13. Olshausen, B. & Millman, K. Learning sparse codes with a mixture-of-Gaussians prior. Advances in Neural Information Processing 12 (2000).

14. Mikami, A., Newsome, W. & Wurtz, R. Motion selectivity in macaque visual cortex. II. Spatiotemporal range of directional interactions in MT and V1. Journal of Neurophysiology 55, 1328–1339 (1986).

15. Ungerleider, L. G. & Haxby, J. V. ‘what’and ‘where’in the human brain. Current opinion in neurobiology 4, 157–165 (1994).

16. Young, M. P. et al. Objective analysis of the topological organization of the primate cortical visual system. Nature 358, 152–155 (1992).

17. Felleman, D. J. & Van Essen, D. C. Distributed hierarchical processing in the primate cerebral cortex. Cerebral cortex 1, 1–47 (1991).

18. Livingstone, M. Mechanisms of direction selectivity in macaque V1. Neuron 20, 509–526 (1998).

19. Fried, S., Munch, T. & Werblin, F. Directional selectivity is formed at multiple levels by laterally offset inhibition in the rabbit retina. Neuron 46, 117–127 (2005).

20. Li, Y., FitzPatrick, D. & White, L. The development of direction selectivity in ferret visual cortex requires early visual experience. Nature Neuroscience 9, 676–681 (2006).

21. Smith, G. B. et al. The development of cortical circuits for motion discrimination. Nature neuroscience (2015).

22. Xu, S., Jiang, W., Poo, M. & Dan, Y. Activity recall in a visual cortical ensemble. Nature Neuroscience 15, 449–455 (2012).

23. Ringach, D. L. Spatial structure and symmetry of simple-cell receptive fields in macaque primary visual cortex. Journal of neurophysiology 88, 455–463 (2002).

24. Reid, R. C., Soodak, R. & Shapley, R. Directional selectivity and spatiotemporal structure of receptive fields of simple cells in cat striate cortex. Journal of Neurophysiology 66, 505–529 (1991).

25. Adelson, E. H. & Bergen, J. R. Spatiotemporal energy models for the perception of motion. JOSA A 2, 284–299 (1985).

26. Reíd, R. C., Soodak, R. E. & Shapley, R. M. Linear mechanisms of directional selectivity in simple cells of cat striate cortex. Proceedings of the National Academy of Sciences 84, 8740–8744 (1987).

27. Linden, J. F., Liu, R. C., Sahani, M., Schreiner, C. E. & Merzenich, M. M. Spectrotemporal structure of receptive fields in areas ai and aaf of mouse auditory cortex. Journal of Neurophysiology 90, 2660–2675 (2003).

28. Geiger, J. R., Lübke, J., Roth, A., Frotscher, M. & Jonas, P. Submillisecond ampa receptor-mediated signaling at a principal neuron–interneuron synapse. Neuron 18, 1009–1023 (1997).

29. Douglas, R., Koch, C., Mahowald, M., Martin, K. & Suarez, H. Recurrent excitation in neocortical circuits. Science 269, 981–985 (1995).

30. Mastronarde, D. N. Two classes of single-input x-cells in cat lateral geniculate nucleus. i. receptive-field properties and classification of cells. Journal of Neurophysiology 57, 357–380 (1987).

31. Vigeland, L. E., Contreras, D. & Palmer, L. A. Synaptic mechanisms of temporal diversity in the lateral geniculate nucleus of the thalamus. The Journal of Neuroscience 33, 1887–1896 (2013).

32. Van Essen, D. C., Anderson, C. H. & Felleman, D. J. Information processing in the primate visual system: an integrated systems perspective. Science 255, 419–423 (1992).

33. Werbos, P. J. Backpropagation through time: what it does and how to do it. Proceedings of the IEEE 78, 1550–1560 (1990).

34. Orban, G., Kennedy, H. & Bullier, J. Velocity sensitivity and direction selectivity of neurons in areas V1 and V2 of the monkey: influence of eccentricity. Journal of Neurophysiology 56, 462–480 (1986).

35. Simoncelli, E. & Heeger, D. A model of neuronal responses in visual area MT. Vision Research 38, 743–761 (1998).

36. Skaggs, W. & McNaughton, B. Replay of neuronal firing sequences in rat hippocampus during sleep following spatial experience. Journal of Neuroscience 17, 2112–2127 (1997).

37. Abbott, L. & Blum, K. Functional significance of longterm potentiation for sequence learning and prediction. Cerebral Cortex 6, 406–416 (1996).

38. Rao, R. & Sejnowski, T. Predictive sequence learning in recurrent neocortical circuits. Advances in Neural Information Processing 12, 164–170 (2000).

39. Dayan, P. & Abbott, L. Theoretical Neuroscience (The MIT Press, 2001).

40. Barber, D. Learning in spiking neural assemblies. Advances in Neural Information Processing 15 (2002).

41. Brea, J., Senn, W. & Pfister, J. Sequence learning with hidden units in spiking neural networks. Advances in Neural Information Processing 24 (2011).

42. Rao, R. & DH, B. Predictive coding in the visual cortex: a functional interpretation of some extra-classical receptive-field effects. Nature Neuroscience 2, 79–87 (1999).

43. Friston, K. A theory of cortical responses. Phil. Trans. R. Soc.B 360, 815–836 (1999).

44. Olshausen, B. & Field, D. Emergence of simple-cell receptive field properties by learning a sparse code for natural images. Nature 381, 607–609 (1996).

45. Rehn, M. & Sommer, F. T. A network that uses few active neurones to code visual input predicts the diverse shapes of cortical receptive fields. Journal of computational neuroscience 22, 135–146 (2007).

46. Wiskott, L. & Sejnowski, T. Slow feature analysis: Unsupervised learning of invariances. Neural Computation 14, 715–770 (2002).

47. Hubel, D. H. & Wiesel, T. N. Receptive fields of cells in striate cortex of very young, visually inexperienced kittens. J. neurophysiol 26, 994–1002 (1963).

48. Blakemore, C. & Cooper, G. F. Development of the brain depends on the visual environment. Nature 477–478 (1970).

49. Hirsch, H. V. & Spinelli, D. Visual experience modifies distribution of horizontally and vertically oriented receptive fields in cats. Science 168, 869–871 (1970).

50. Miller, K. D., Erwin, E. & Kayser, A. Is the development of orientation selectivity instructed by activity? Journal of neurobiology 41, 44–57 (1999).

51. Sengpiel, F., Stawinski, P. & Bonhoeffer, T. Influence of experience on orientation maps in cat visual cortex. Nature neuroscience 2, 727–732 (1999).

52. Tanaka, S., Ribot, J., Imamura, K. & Tani, T. Orientation-restricted continuous visual exposure induces marked reorganization of orientation maps in early life. Neuroimage 30, 462–477 (2006).

53. Berkes, P., Turner, R. & Sahani, M. A structured model of video produces primary visual cortical organisation. PLoS Computational Biology 5 (2009).

54. Ko, H. et al. Functional specificity of local synaptic connections in neocortical networks. Nature 473, 87–91 (2011).

55. Ko, H. et al. The emergence of functional microcircuits in visual cortex. Nature 496, 96–100 (2013).

56. Minka, T. Expectation propagation for approximate Bayesian inference. UAI’01 Proceedings of the Seventeenth conference on Uncertainty in artificial intelligence 362–369 (2001).

57. Doucet, A., Freitas, N., Murphy, K. & Russell, S. Rao-blackwellised particle filtering for dynamic Bayesian networks. UAI’00 Proceedings of the Sixteenth conference on Uncertainty in artificial intelligence 176–183 (2000).

58. Goodfellow, I., Courville, A. & Bengio, Y. Spike-and-slab sparse coding for unsupervised feature discovery. arXiv 1201.3382v2 (2012).

59. Mallat, S. & Zhang, Z. Matching pursuits with time-frequency dictionaries. IEEE Transactions on Signal Processing 41, 3397–3415 (1993).

